# Enhanced capillary delivery with nanobubble-mediated blood-brain barrier opening and advanced high resolution vascular segmentation

**DOI:** 10.1101/2023.12.14.571641

**Authors:** Roni Gattegno, Lilach Arbel, Noa Riess, Sharon Katz, Tali Ilovitsh

## Abstract

Overcoming the blood-brain barrier (BBB) is essential to enhance brain therapy. Here, we utilized nanoscale nanobubbles with focused ultrasound for targeted and improved BBB opening in mice. A microscopy method assessed BBB opening at a single blood vessel resolution employing a dual-dye labeling technique using green fluorescent molecules to label blood vessels and Evans blue brain-permeable dye for quantifying BBB extravasation. A UNET-based deep learning architecture enabled blood vessels segmentation, delivering comparable accuracy to manual segmentation with a significant time reduction. Segmentation outcomes were applied to the Evans blue channel to quantify extravasation of each blood vessel. Results were compared to microbubble-mediated BBB opening, where reduced extravasation was observed in capillaries with 2-6μm diameter. In comparison, nanobubbles yield an improved opening in these capillaries, and equivalent efficacy to that of microbubbles in larger vessels. These results indicate the potential of nanobubbles to serve as enhanced agents for BBB opening, amplifying bioeffects in capillaries while preserving comparable opening in larger vessels.

## Introduction

The blood brain barrier (BBB) is a protective barrier formed by specialized endothelial cells that line the blood vessels in the brain that tightly regulates the passage of substances between the bloodstream and the brain tissue^1^. While the BBB is crucial for maintaining the microenvironment of the brain and protecting it from potentially harmful substances, it also poses a challenge for delivering therapeutic agents, including drugs^2^. Microbubbles (MBs), in combination with low frequency focused ultrasound (FUS), have been at the forefront of therapeutic efforts to open the BBB for many brain related conditions^3–5^. Currently, FUS-mediated BBBD clinical trials are ongoing for brain diseases such as recurrent glioblastoma and Alzheimer’s disease, showing the promise of this technique^6,7^. Most of these trials use low frequency FUS, on the order of 250 kHz, to improve skull penetration in a noninvasive manner. When FUS is applied to gas filled MBs with a diameter of 1.5-4 μm, it causes them to expand and contract repeatedly. As this process occurs within blood vessels in the brain, the BBB can open safely and transiently to facilitate drug delivery to targeted region. Therefore, MB oscillations play a key role in successful BBBD. We recently showed that molecule extravasation following MB-mediated BBBD is reduced in capillaries^8^. It is likely due to a combination of effects, including constrained oscillations alongside smaller numbers of MBs within these capillaries, compared to larger arteries^9–11^. This scenario becomes more complex when considering the inherent heterogeneous diameters of brain vasculature, ranging from capillaries to arteries, which leads to variations in BBBD as a function of blood vessel diameter and can potentially inhibit desired therapeutic results in smaller vessels^12,13^.

To bridge these gaps, we propose to use nanobubbles (NBs) for BBBD. Their smaller size on the range of 200 nm, could facilitate their entry into capillaries, increasing their numbers in smaller vessels. NBs were used for ultrasound contrast and molecular imaging^14,15^ Cancer imaging ^16,17^, sonoporation^18^, gene delivery^19,20^, Molecular imaging, noninvasive cancer therapy^21,22^, and BBBD^23–27^. Prior studies on NB-mediated BBBD primarily focused on establishing NBs for this purpose^23–27^. Here, our objective was to investigate BBBD at the resolution of single blood vessels, specifically targeting extravasation patterns in capillaries and comparing the results to those obtained with MBs. We were particularly interested in the context of low-frequency applications at 250 kHz, which is well-suited for brain therapy. At this frequency, we have observed high-amplitude MB oscillations induced by the Blake threshold effect^28–31^, and we have successfully applied this effect to enhance the response of NBs for cancerous tumor mechanotherapy^21^. In this study, we propose that these amplified oscillations, combined with the reduced size of NBs, will enhance their bioeffects in capillaries by increasing their presence within these vessels and allowing them to oscillate freely, unlike MBs, which may face constraints in smaller vessels. To evaluate the success of BBBD and compare the performance of NB to that of MB, BBBD extravasation assessment was used.

The common methods for this task include two-photon microscopy and magnetic resonance imaging (MRI)^32,33^. Two-photon microscopy offers in-depth, high-resolution images, and is known for its efficacy in visualizing complex biological processes in vivo. However, while two-photon microscopy provides detailed imagery, its penetration depth is limited, and it includes a tradeoff between spatial resolution and acquisition time, that typically prevent the imaging of capillaries below 10 µm. MRI can provide a qualitative extravasation assessment yet cannot be used for single blood vessel assessment.

Here, we adopted a single blood vessel BBBD analysis technique that enabled the quantification of EB extravasation in blood vessels smaller than 10 µm at a resolution of 1 µm^8^. This method was carried on full brain slices and was not subjected to the penetration depth limitations that exist with two-photon microscopy. The method used a dual staining technique that involved the injection of two fluorescent dyes; the first was Evans-blue (EB) that served as a marker for BBB extravasation, and the second was a large, FITC-labeled dextran (FITC), that served as a marker of the blood vessels. Standard fluorescence microscopy of brain slices was then used, and an automated image processing pipeline was developed to assess EB extravasation at a single blood vessel resolution and compare the results to MB-mediated BBBD (Fig. 1). The capture imaged from the brains were segmented according to the FITC channel and the segmentation was then applied to the EB channel to quantify the BBBD. The automated blood vessel segmentation technique employed a machine learning algorithm to extract the geometrical features of the blood vessels. In general, blood vessel segmentation serves as a support tool with diverse clinical domains, notably in neurosurgery and laryngology^34,35^. Distinguishing vessels from adjacent tissues, can aid in diagnosis, treatment planning, and the assessment of clinical outcomes^34,36^. While traditional manual segmentation, remains essential, it can be labor-intensive and error-prone^36,37^. The evolution of automatic segmentation techniques allows for an in-depth exploration of the morphology, orientation, and distribution of blood vessels^34,36^. This segmentation is essentially a classification challenge, ensuring each pixel’s categorization as either a vessel or background^34,37^. The efficacy of deep neural networks, such as CNN, 3D-CNN, and U-NET, in these tasks is notable^36^. Here, we chose to use a well-established U-NET architecture to enhance the speed and accuracy of blood vessel segmentation. Integrating machine learning based segmentation techniques with BBBD assessment constitute a robust platform for evaluating the impact of microvasculature diameter on BBBD and for comparing the effectiveness of NB and MB in brain therapy applications.

**Figure 1.**
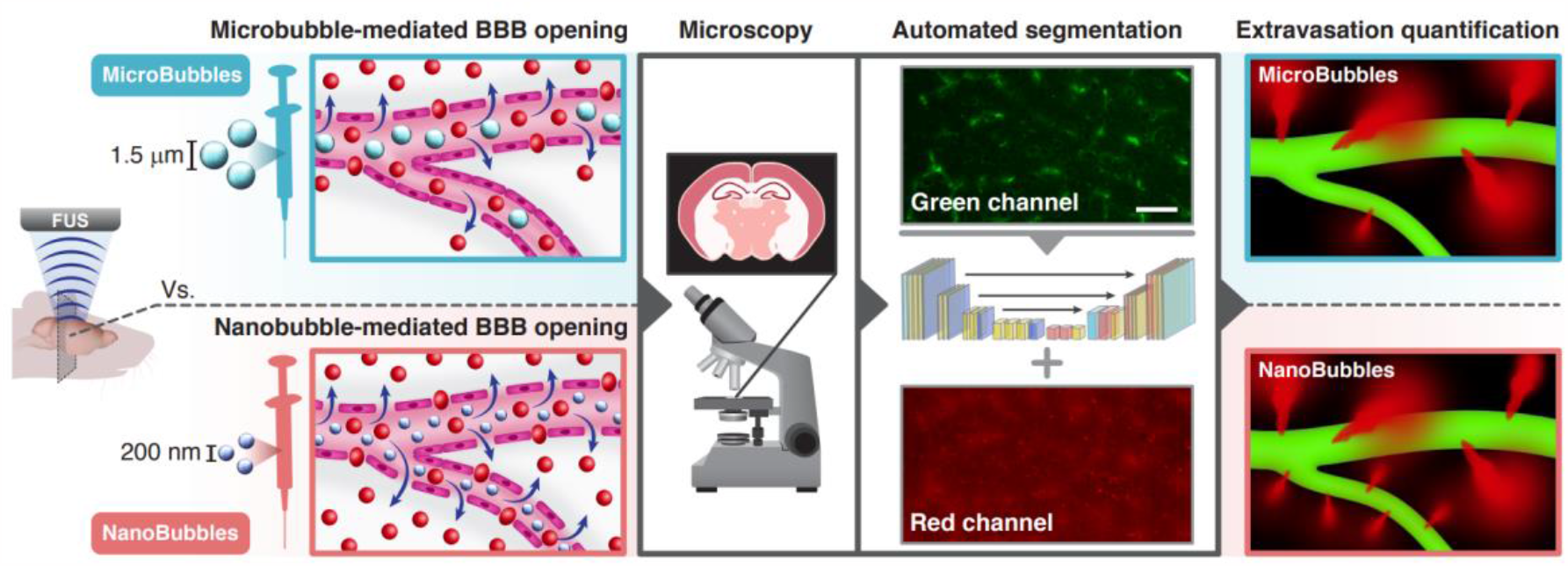
Extravasation assessment of bubble-mediated BBB opening. BBB opening was induced using MB (∼1.5 µm) or NB (∼200 nm). Following BBBD, mice were injected with two dyes: a large FITC dye to mark the blood vessels and EB that can extravasate into BBBD regions. Fluorescence microscopy imaged brain slices, and an automated segmentation process employing a neural network classifier segmented the blood vessels. Lastly, EB extravasation as a function of blood vessel diameter was assessed, comparing MB and NB BBBD.

## Results

### NB-mediated BBBD in mice

MBs and NBs with a diameter of 1.18 µm and 160 nm, respectively, were fabricated. First, the safe range of BBBD was assessed for PNPs in the range of 150-300kPa and a center frequency of 250 kHz. The range between 175-250 kPa yielded BBBD without inducing any observable functional damage, or microhemorrhage. As such, this safe range was consistently employed in all subsequent experiments. Importantly, this is the same range of PNPs that was used for MB-mediated BBBD. Microscopy imaging confirmed BBB opening and assessed the EB extravasation as a function of the blood vessel diameter. A 2 MDa FITC injection was administered to label the blood vessels in the green channel, while the red channel displayed the EB-injected dye. In control mice, the green and red channels overlapped, and there was no EB leakage since the BBB was intact (Fig. 2A-C). In the treated group, EB extravasation was observed mainly in the cortex region (Fig 2D-F). To quantify BBBD at a single blood vessel resolution, 20x magnified images were captured at the region of BBBD for the control mice, the MB + FUS, and NB + FUS groups (Fig. 3). As observed in Fig. 2A-C, the green and red channels overlapped in control with an intact BBB (Fig. 3A). When qualitatively comparing images in the MB + FUS (Fig. 3B) vs. NB + FUS (Fig. 3C), the amount of red extravasation in the frames is higher, implying that more dye leaked. Next, the amount of EB extravasation was quantified as a function of blood vessel diameter using a post processing pipeline (Fig. 3D-I). The green channel (Fig. 3D) was employed for blood vessel segmentation and shape delineation through neural network automated segmentation, with a comparison to manual processing (Fig. 3E,F). The segmented vessels were then integrated with the red channel, facilitating the computation of extravasated EB around each blood vessel (Fig. 3G-I).

**Figure 2.**
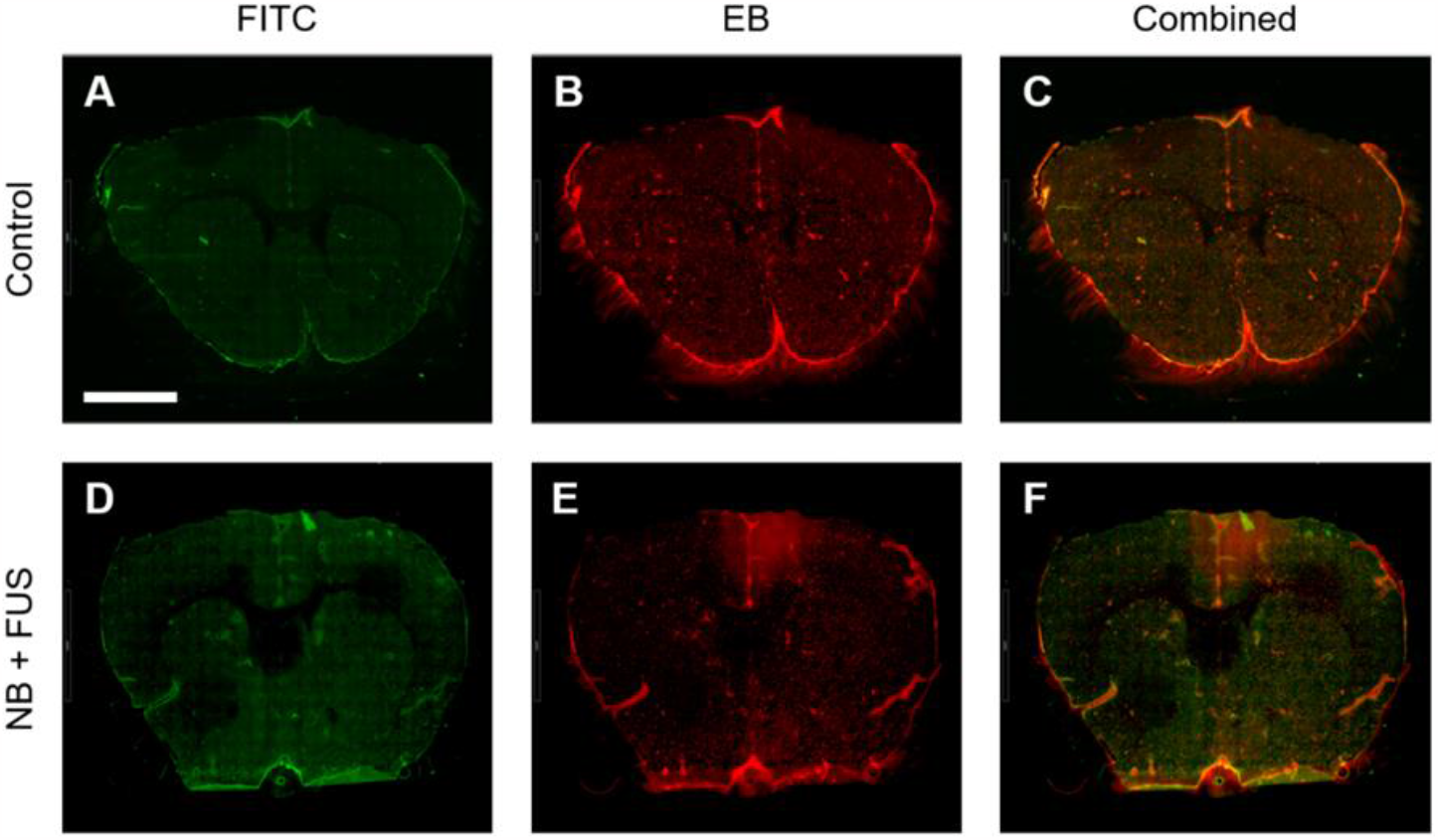
Microscopy images of brain slices. Fluorescent microscopy images of coronal slice from (A)-(C) control group, and (D)-(F) NB group. (A), (D) Green FITC fluorescent dye highlights the blood vessels. (B), (E) Evans blue extravasation as detected in the red channel. (C) is an overlay of (A) and(B). (F) is an overlay of (D) and (E). The images were acquired with 20x objective lens. Scale bar: 2 mm.

**Figure 3.**
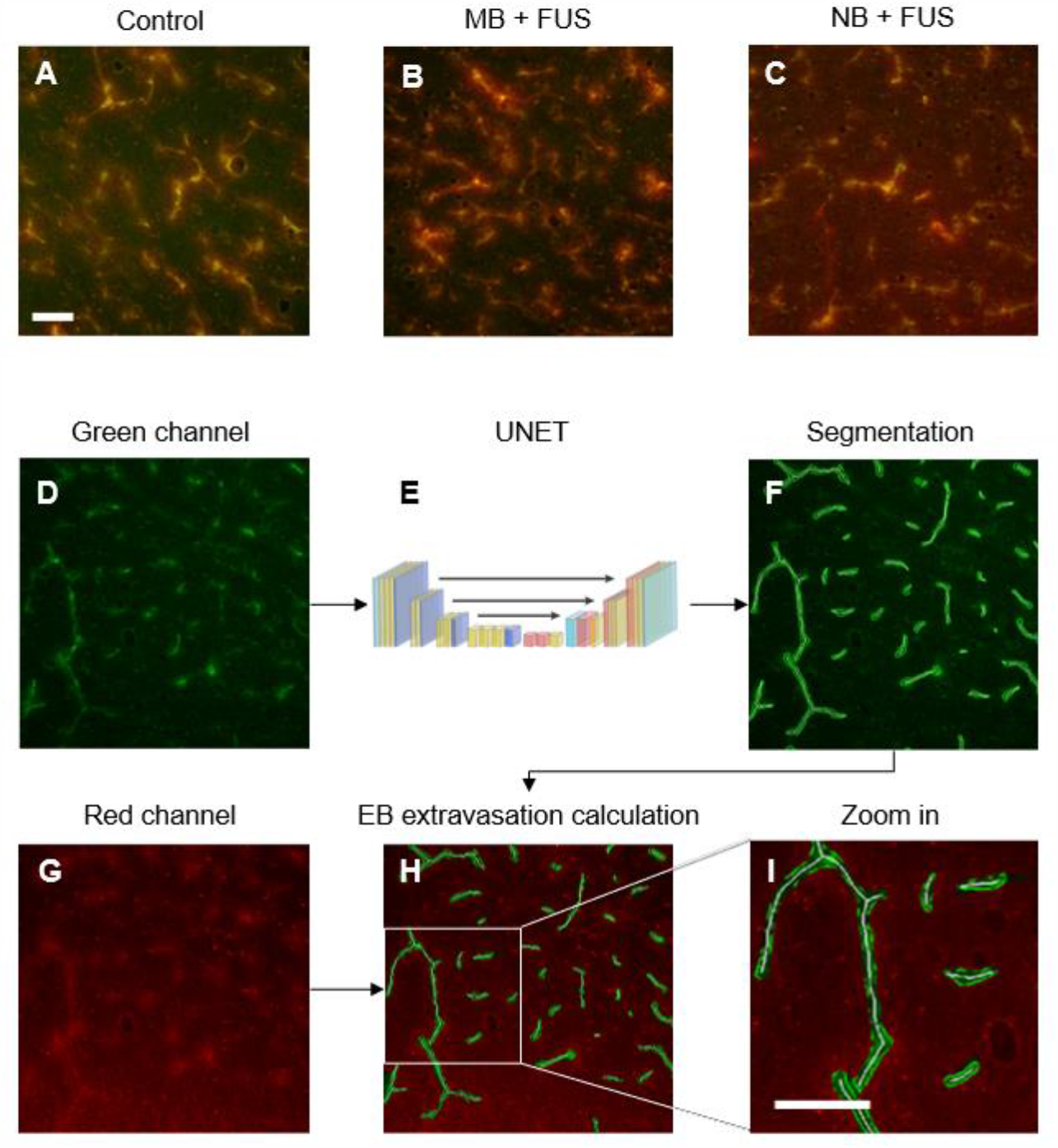
Image processing pipeline of microscopy images. Microscopy images of brain slices from (A) control group, (B) MB group and (C) NB group. The images were captured from the cortex using a 20x objective lens. (D-I) Image processing procedure: the green channel (FITC-Dextran, D), indicating blood vessels undergoes either (E) UNET-automated segmentation or (F) manual segmentation to extract geometrical features. This segmentation is then integrated into the red channel (G) that signifies EB extravasation, enabling the quantification of EB intensity around each vessel (H). Subfigure (I) provides a magnified view of the red channel with overlaying segmentation. Scale bars are 50 μm in all subfigures.

### UNET assisted blood vessel segmentation

The blood vessel segmentation was then applied to the red channel to evaluate the amount of EB Automated blood vessel segmentation utilized a U-NET architecture^38^. The dataset contained 368 manually segmented frames of 21,491 blood vessels. The training set, comprising 85% of the total frames, underwent training across more than 100 epochs with a batch size of 8. Initially, the efficacy of the model was evaluated by comparing the results with manually segmented images, which served as the ground truth. The performance of the classifier was tested on different images from various brain regions, showing the binary prediction of the model against the manually segmented ground truth and highlighting the discrepancies in the difference image (Fig. 4A). An AUC of 0.9918, an F1 score of 0.743, and a recall rate of 0.9065 were achieved by the model, as derived from the confusion matrix (Fig. 4B). Subsequently, EB extravasation around manually and automated segmented blood vessel from the NB group were analyzed (Fig. 4C). In the analysis, 2,179 blood vessels from the manually segmented set and 1,151 from the combined automated and manual refinement set were included. No significant differences across all vessel size categories were indicated by a multi-comparison two-way ANOVA test (p value >0.05).

**Figure 4.**
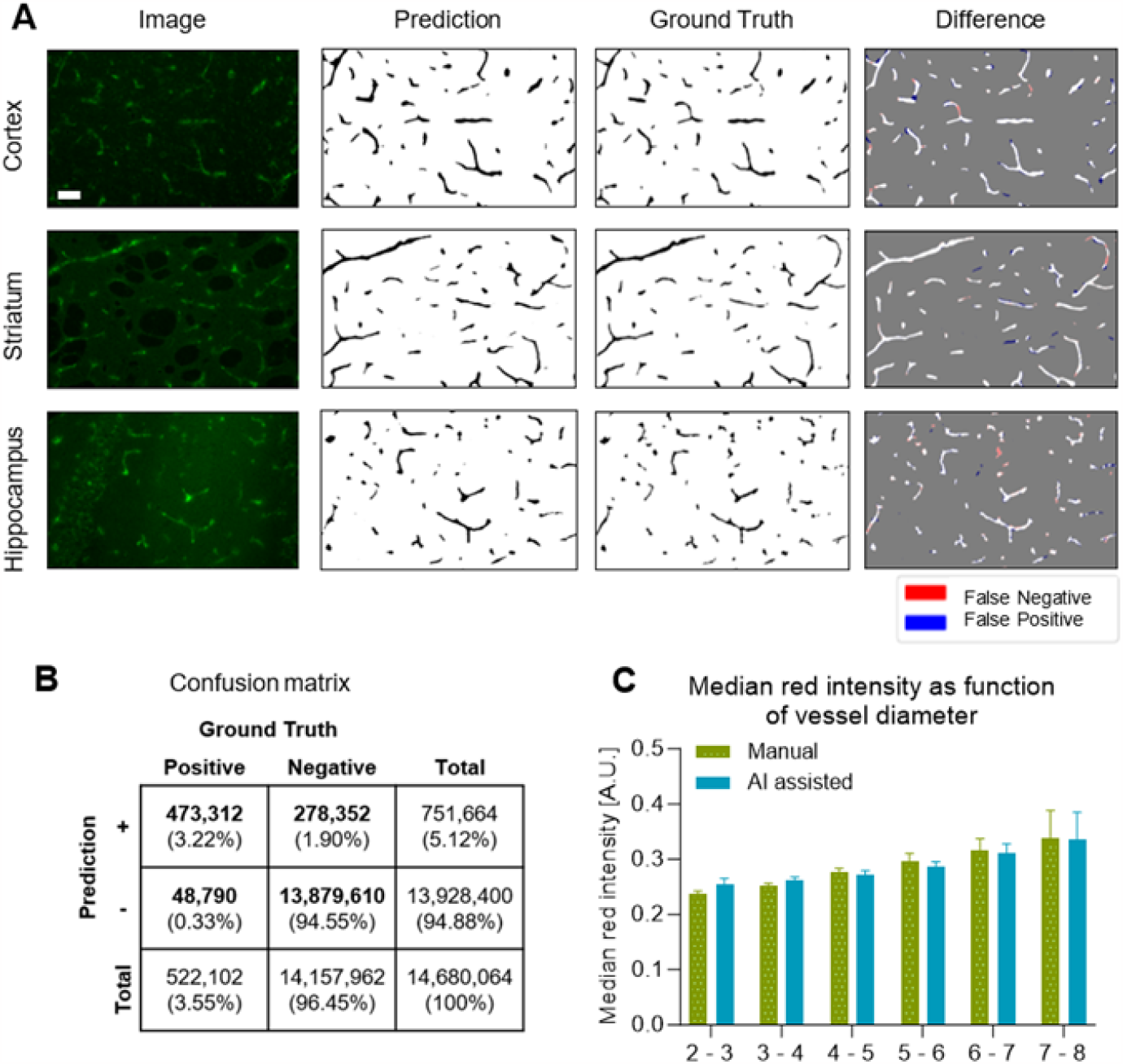
Neural-network assisted automated blood vessel segmentation. (A) Performance of the classifier on different images from different brain regions. The binary prediction of the model, the manual segmentation as the ground truth and the difference is shown for each image. In the difference image, false negative predictions marked in red and false positive prediction marked in blue. (B) Confusion matrix representing the performance of the automated segmentation process. (C) EB extravasation observed during NB-mediated BBBD in relation to microvasculature diameter. Comparisons were made between manually segmented images and those segmented via AI and manually revised. In total, the datasets comprised n = 2179 blood vessels for manual segmentation and n = 1151 for AI-assisted segmentation. Two-way ANOVA test was applied. No significant difference in any of the size groups (p value >0.05).

### EB extravasation assessment

The blood vessel segmentation was then applied to the red channel to evaluate the amount of EB leakage around each blood vessel in the three groups that were tested (control, MB + FUS, NB + FUS). The extravasation around each blood vessel was evaluated as a function of the distance from the vessel wall spanning 0.3-0.9 μm, and as a function of the perivascular area spanning 1.5-3 μm. It was observed that varying these parameters impacted all groups in a similar manner, not significantly altering the comparative results (Fig. S1). Therefore, we chose parameters that were previously used^8^: A distance of 2 pixels (0.58 μm width) from the vessel wall and a perivascular area extending 10 pixels (2.93 μm width(. An analysis of 4,500 blood vessels from the NB group, 2,877 from the MB group, and 3,403 from the control group was conducted. The blood vessel diameter, on average, across the three groups was comparable, with values of 4.11±1.22, 4.08±1.38, and 3.92±1.37 respectively (mean ± SD, Fig. 5A). No significant difference was found between the variance in the diameter histograms (Ordinary one-way ANOVA test, p>0.05).

**Figure 5.**
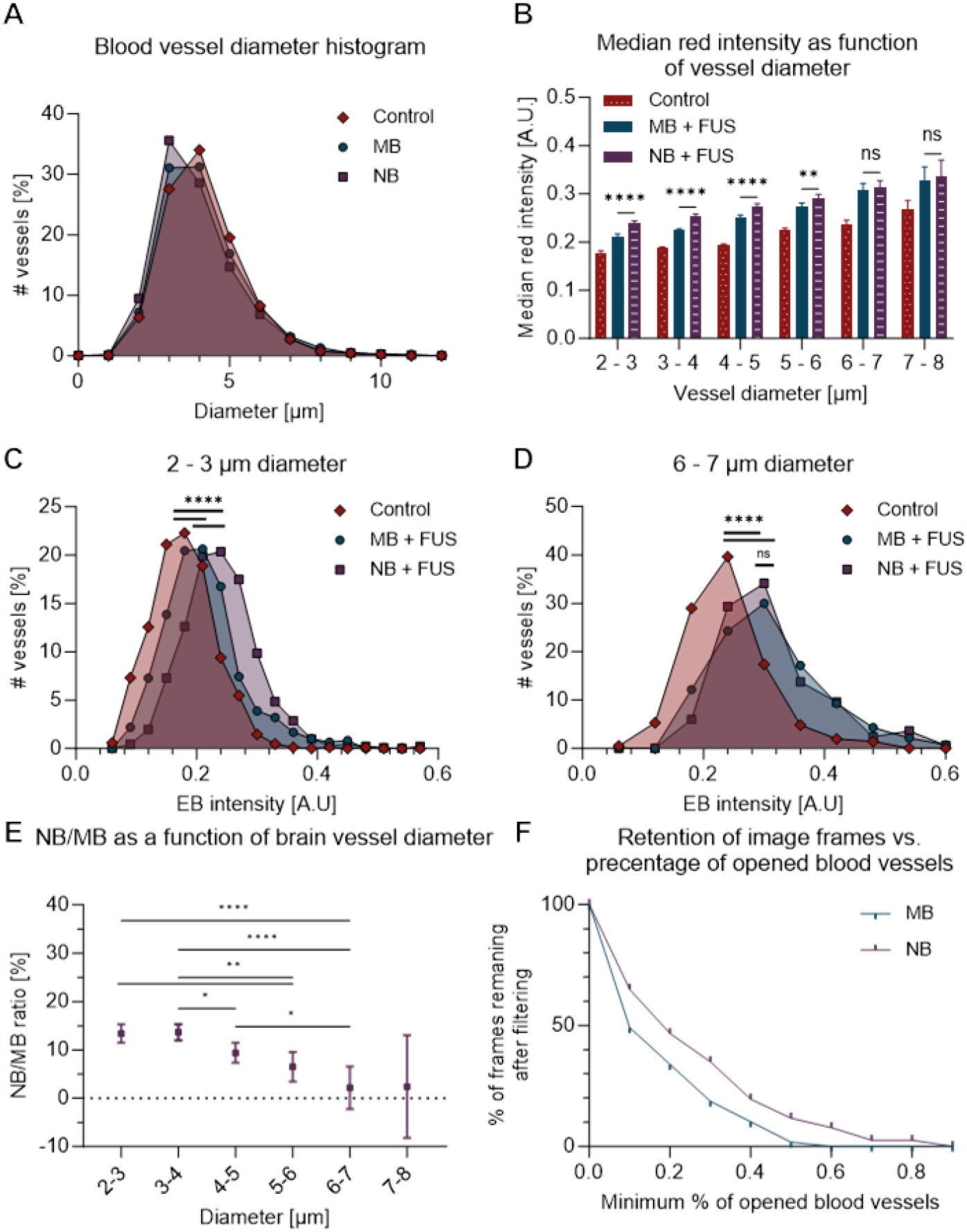
Enhanced EB extravasation in small blood vessels using NB-mediated BBBD. (A) Blood vessels diameter histogram (total of n = 4500, n = 2877 and n = 3403 blood vessels for the control, MB + FUS and NB + FUS groups, respectively). (B) EB extravasation quantification as a function of blood vessel diameter. The graph is plotted as mean + 95% CI. Two-way ANOVA with Tukey correction. The p values shown are between MB and NB; ^*^p < 0.05, ^**^ p < 0.01, ^***^p < 0.001, ^****^p < 0.0001. All p values between MB and control and between NB and control are less than 0.0001. (C and D) The EB intensity histogram for blood vessels with diameters of (C) 2 – 3 μm and (D) 6 – 7 μm. (E) NB/MB ratio as a function of brain vessel diameter. One-way ordinary ANOVA test with Tukey correction. P values as in C and D. (F) Retention of Image frame vs. percentage of opened blood vessels (total of n = 59 and n = 77 frames for MB + FUS and NB + FUS groups, respectively).

Next, EB extravasation as a function of blood vessel diameter was assessed. Within each of the three groups, EB extravasation increased with blood vessel diameter (2-way ANOVA test, p value <0.0001). Within every diameter-size category that was examined (ranging from 2-8 μm), statistically significant differences were detected between the MB + FUS and control groups, as well as between the NB + FUS and control groups (p value< 0.0001, Fig. 5B). In the larger blood vessels tested (specifically in the 6-7 μm and 7-8 μm ranges), comparable results between the MB + FUS and NB + FUS groups were observed, with no significant variance being detected (p value>0.05). However, in the narrower 2-6 μm vessels, a higher EB extravasation intensity was exhibited by the NB + FUS group. This can also be seen when looking at individual EB extravasation histograms for specific diameters. For a diameter of 2-3 μm, there is a separation between the control, MB + FUS and NB + FUS groups, with the highest average EB intensity individual EB extravasation histograms for specific diameters. For a diameter of 2-3 μm, there is a separation between the control, MB + FUS and NB + FUS groups, with the highest average EB intensity observed in the NB + FUS group (Fig. 5C). For a diameter of 6-7 μm, the MB + FUS and NB + FUS group become similar yet separated from the control group (Fig. 5D). A comparative analysis of the normalized EB extravasation ratio between the MB + FUS and NB + FUS groups, which was calculated based on Eq. 5, indicated a 13% improvement for the NB + FUS group in blood vessels of 2-4 μm. The improvement decreased as the blood vessel size increased, indicating that BBB in larger vessels was not affected, yet only improved in small blood vessels (Fig. 5E). To quantify the robustness of BBBD, a threshold was set to determine the number of frames with strong BBBD. This threshold, calculated for each diameter-size group, was defined as two standard deviations above the average of the control group within the same diameter-size category. Subsequently, the count of opened blood vessels per frame was calculated, and the ratio of opened vessels to total vessels was computed. For filtration thresholds ranging from 0.1 to 0.9, frames with ratios below the threshold were excluded, and the remaining frames were calculated (Fig. 5F). The NB + FUS group consistently retained a greater number of frames compared to the MB + FUS group, irrespective of the chosen filtration threshold, which suggests that BBBD was more pronounced in the NB + FUS group.

## Discussion

This study investigates the utilization of NBs as improved theranostic agents for brain therapy applications, aiming to optimize their performance within capillaries. The safe range of PNPs for BBBD was identified as 175-250 kPa for a center frequency of 250 kHz, both for the MB and NB groups. This result is intriguing considering the significant size difference between these agents, as it might be expected that NBs would necessitate higher PNPs. One possible explanation for the strong effect in the NB group is that exciting NBs within the same frequency range as MBs could significantly enhance their oscillations, as the Blake threshold applies to gas-bubble excitation well below their resonance frequency. The use of NBs could improve both their penetration into smaller capillaries and enhance therapeutic applications through improved oscillations. It was shown that for MB-mediated BBBD, EB extravasation increases with blood vessel diameter^8^. Nevertheless, extravasation in capillaries with diameter of 2-6 µm is important both because they constitute the majority of brain blood vessel in mice, and because capillaries play a key role in the pathophysiology of multiple brain diseases such as small vessel disease, a leading cause of stroke and cognitive decline in the elderly, and Alzheimer’s disease, the most prevalent elderly dementia^39–41^. Reducing the size of MBs to the nanoscale was shown here to improve BBBD within small capillaries, without affecting the performance in the larger vessels.

To probe BBBD at a single blood vessel resolution, a dual dye labeling technique following BBBD using bubble and US was used. Large FITC-dextran molecules were injected to label the blood vessels, and EB dye was used to assess BBB extravasation. Brain slices were imaged via fluorescence microscopy and an image processing method was employed to evaluate the extravasation as a function of vasculature diameter, while comparing the results of MB and NB-mediated BBBD. The green channel was used for blood vessel segmentation. Here, unlike previous work^8^, which used only manual segmentation of the blood vessels, we implemented a machine learning based algorithm for this task. Our approach was based on the U-NET architecture, an encoder-decoder architecture tailored for pixel-level segmentation in medical imagery^42^. When the segmentation performance was tested, comparing the automated segmentation to the ground truth manually segmented data, AUC of 0.9918, F1 score of 0.743, and a recall rate of 0.9065 were achieved. Comparing EB extravasation between automated-assisted segmentation and manually segmented frames, we find no significant difference in the EB extravasation between the two segmented data (p = 0.9767). The use of machine learning allowed us to merge automation with human oversight for optimal outcomes, streamline image analysis process and analyze more images from each brain. Manual effort and time were reduced by approximately 90% using this technique.

The segmentation process outputs a mask, which was applied to the red channel to calculate the EB extravasation around each blood vessel. In direct comparison to MB-mediated methods, NBs showcased superiority, particularly evident in a 13% improvement of EB extravasation in smaller blood vessels (2-6 μm), while differences in larger vessels remained statistically insignificant. Furthermore, we showed that BBBD is more uniform in the NBs group by setting a diameter-based threshold to compute the fraction of strong BBBD vessels in each frame. Frames were filtered using thresholds from 0.1 to 0.9, keeping only those with an opened-to-total vessel ratio above the set threshold. Across all thresholds, the NB group consistently had a higher frame retention compared to the MB group with max improvement of 15.8% for threshold of 0.1.

Our findings underline the potential of NBs as a promising tool in BBBD. Their superior performance in smaller vessels, combined with equivalent efficacy to that of MBs in larger vessels, suggests that NBs can be used as targeted approach for refined brain therapeutic applications. Future techniques could utilize the NBs ability to accumulate within leaky tumors due to their reduced size and incorporated in brain therapy applications^16,17^.

Here we focused on assessing BBBD mainly in the cortex. It is possible that BBBD is affected by additional parameters such as blood vessels density and morphological features such as length and bifurcation, that are region dependent. Therefore, in future studies, a region-based approach, encompassing diverse brain regions, would be beneficial to fully understand the differential effects across the brain. Moreover, blood vessels are inherently three-dimensional, while our microscopy images are two-dimensional, which might affect the classification and extravasation assessment accuracy. It will be possible to use brain clearing techniques combined with 3D microscopy to visualize full brain architecture and quantify extravasation. Additionally, our analysis was limited to coronal slices. Incorporating sagittal and horizontal slices in future research will provide alternative perspectives on the brain.

An EB dye was used as a marker to assess BBBD. EB is a relatively small molecule (∼1 kDa), and its extravasation patterns might not accurately represent those of larger molecules or particles. This is particularly relevant in the context of neurotherapeutics or diagnostic agents that are often larger in size than EB. Therefore, the results obtained using EB might not fully capture the dynamics or extent of BBBD for these larger entities, suggesting a need for future studies to explore BBBD with markers of varying molecular sizes. In addition, BBBD utilized NBs. Their concentration and size distribution were measured using a particle sizing system (AccuSizer FXNano). This system cannot distinguish between liposomes and NBs. This overlap in measurement implies that the actual count of NBs could have been lower than what was reported.

For the automated image processing, a U-NET architecture tailored for medical image segmentation was used^38^. Following the introduction of U-Net, subsequent advancements in the field have emerged, exceeding its performance^36,43^. It is important to note that the application of U-NET in this study was mainly as an assist tool. Although it dramatically reduced time, manual refinement was still required. Considering the broad spectrum of available machine learning models, it is possible that the optimization of parameters or the adoption of other models, like CNN^37^, might replace the manual process entirely, promising a more robust solution. Moreover, the presence of manual interventions (either as a refining step post-automated segmentation or as entirely manual segmentation) could potentially introduce subjective biases, a challenge that might be overcome with a fully automated approach. Lastly, this study primarily centers on the immediate and short-term consequences of post-treatment. Given the potential therapeutic applications of NB-mediated BBBD, future research must delve deeper into long-term effects to fully understand its implications.

## Materials and Methods

### Experimental setup

The experimental setup consisted of a spherical single-element focused ultrasound (FUS) transducer (H115, Sonic Concepts, Bothell, WA, USA) operating at a center frequency of 250 kHz, as described in previous works^44–46^. The transducer has a focal distance of 45 mm and an aperture diameter of 64 mm (Fig. 6). The measured focal spot of the transducer was 8 mm laterally and 50 mm axially, which was deemed suitable for inducing BBBD in large brain regions. Positioned at the bottom of a water tank and oriented upwards (Fig. 6B), the transducer was activated using a transducer power output system (TPO-200, Sonic Concepts, Bothell, WA, USA). Calibration of the peak negative pressure (PNP) at the focal spot used a needle hydrophone (Model NH0500, Precision Acoustics, UK). An agarose phantom was placed at the transducer focal spot, between the water tank and the mice cranial region (Fig. 6A). The agarose phantom was fabricated by mixing 1.5% agarose powder (Product Code: A10752, Alfa Caesar, MA, USA) with deionized water. The resulting solution was heated until complete dissolution of the powder, after which it was poured into a custom-designed mold. For immobilization of the mice and administration of anesthesia during the treatment procedure, a custom-made head holder was positioned above the water tank. This head holder was designed to securely hold the heads of the mice in place throughout the course of the experiment (Fig. 6A).

**Figure 6.**
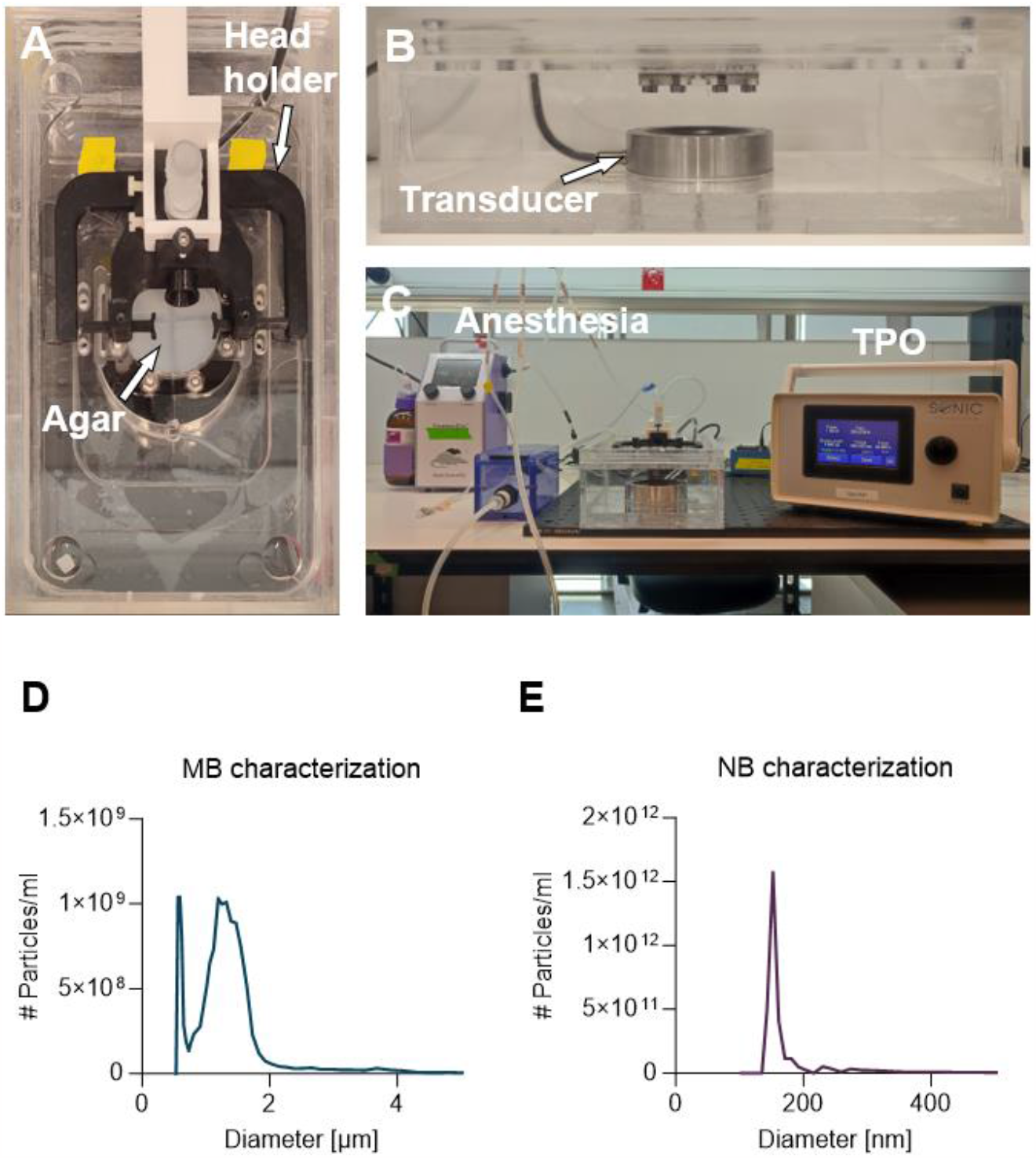
Experimental setup and bubbles characterization. (A) Top view of the setup including a 250 kHz single element transducer at the bottom of a water tank. A head holder located on top of the upper plate is used to position the mice head at the FUS focal spot, and supply anesthesia. For impedance matching, an agarose gel-pad is placed below the mouse head. (B) Side view of the setup including the water tank and the transducer contained within as described in previous work^47^. (C) Overall view of the setup containing the anesthesia machine and the transducer power output used to operate the transducer (TPO). (D) MB concentration as a function of diameter. The median diameter is 1.18 μm.(E) NB concentration as a function of diameter. The median diameter is 160 nm.

### Microbubble preparation

MBs were prepared as described in earlier studies^48^. The MBs consisted of a phospholipid shell encompassing a core of perfluorobutane (C4F10) gas. Distearoylphosphatidylcholine (DSPC; 850365C) and 1,2-distearoyl-sn-glycero-3-phosphoethanolamine-N-[methoxy(polyethylene glycol)-2000] (ammonium salt) (DSPE-PEG2K; 880129C) (Sigma Aldrich, St Louis, MO, USA) lipids were combined in a molar ratio of 90:10 and prepared using the thin film hydration method. The lipids were used in the amount of 2.5 mg per 1mL. To form the MBs, a buffer solution consisting of a mixture of glycerol (10%), propylene glycol (10%), and saline (80%) with a pH of 7.4 was added to the lipids, and the mixture was sonicated at 62°C. Subsequently, the MB precursor solution was divided into vials, each containing 1 mL of liquid, and then saturated with perfluorobutane. Before usage, the vials were agitated for 45 seconds in a vial shaker (Bristol-Myers Squibb Medical Imaging Inc., N. Billerica, MA), and a centrifugation step was employed to remove MBs with radii smaller than 0.5 µm. Another size selection process was performed to eliminate MBs with diameters larger than 5 µm, as described in previous works. The size and concentration of the prepared MBs were determined using a particle counter system (AccuSizer® FX-Nano, Particle Sizing Systems, Entegris, MA, USA). The MBs median diameter was 1.18μm (Fig. 6D), and they were used within three hours from the time of their preparation. Notably, the size distribution and concentration of the MBs exhibited a variation of less than 10% between successive measurements.

### Nanobubble preparation

The NBs were prepared as described previously^21,49^. The NBs consisted of a phospholipid shell encompassing a core of octafluoropropane (C3F8) gas. To prepare the phospholipid solution (10 mg mL−1), 1,2-dibehenoyl-sn-glycero-3-phosphocholine (C22, Avanti Polar Lipids Inc., Pelham, AL), 1,2-dipalmitoyl-sn-glycero-3phosphate (DPPA, Corden Pharma, Switzerland), 1,2-dipalmitoyl-sn-glycero-3-phosphoethanolamine (DPPE, Corden Pharma, Switzerland), and 1,2-distearoyl-snglycero-3-phosphoethanolamine-N-[methoxy(polyethylene glycol)-2000] (ammonium salt) (DSPE-mPEG2000, Laysan Lipids, Arab, AL) were dissolved in propylene glycol (PG, Sigma Aldrich, Milwaukee, WI) by heating and sonicating at 80°C. Next, a mixture of glycerol (Gly, AcrosOrganics) and phosphate buffer solution (0.8 mL, Gibco, pH 7.4) preheated to 80 °C, was added to the lipid solution. The resulting solution was sonicated at room temperature for 10 minutes. A 1 mL portion of the solution was then transferred to a 3 mL headspace vial, which was capped with a rubber septum and an aluminum seal. The vial was sealed using a vial crimper. Air was manually removed from the vial with a 30 mL syringe, and octafluoropropane gas was injected to replace the air. Upon use, the NBs were activated by mechanical shaking with a VialMix shaker for 45 seconds. NB were then separated from the mixture of foam and microbubbles through centrifugation at 50 rcf for 5 minutes with the headspace vial inverted. The resulting 100 μL NB solution was withdrawn from a fixed distance of 5 mm from the bottom of the vial using a 21 G needle. NBs were measured using the same particle counter system as the one used for MB measurements. The NBs median diameter was 0.16 μm (Fig. 6E).

### In-vivo BBBD Experiments

In the in-vivo experiments, a total of 30 female C57BL/6 mice (10-12 weeks old, 16-21g, Envigo, Jerusalem, Israel), were used. The mice were divided into three groups: five mice for each treatment group (MB + FUS and NB + FUS) and three mice for the control group. All animal experiments were carried out following the Guide for the Care and Use of Laboratory Animals and were approved by the Institutional Animal Care and Use Committee at Tel-Aviv University. Before the experiments, the mice were anesthetized with 1.2% isoflurane using a low flow vaporizer system (120 ml/min, SomnoFlo, Kent Scientific). The head hair of the mice was removed with hair removal cream, and US gel was applied. The mice were positioned in a supine position on top of an agarose pad, at the focal spot of the FUS setup described previously. BBBD was induced in the treated groups through retro orbital injection of bubbles. The MB treatment group received 2 x 107 MBs in 50 µl of degassed PBS, following a protocol from previous work^8^. The NB treatment group was injected with 0.737 µl to 1 g of animal weight, diluted in a degassed PBS in a1:1 ratio, as described in other works29,43. NBs dosage was chosen to maintain a comparable gas volume to that of the MB treatment^50^. Thirty seconds after the bubble injection, FUS treatment was administered, comprising 1 ms bursts at 250 kHz with a total sonication period of 60 seconds and a PRF of 1 Hz (0.1% duty-cycle). The Sonication PNP was 150-300 kPa and was optimized to induce safe, mild BBBD, visible in fluorescent microscopy, without causing microhemorrhages in histology. Following FUS treatment, the mice were systemically injected with two dyes: 2% Evans Blue dye (EB; E2129, Sigma Aldrich) (4 ml/kg, diluted in PBS), and 6 mg/ml 2000 kDa Fluorescein isothiocyanate–Dextran (FITC-Dextran; 46946, Merck, Kenilworth, NJ, USA) (4 ml/kg, diluted in normal saline). EB circulation time was 18 minutes, as optimized in previous work^8^. FITC circulation time was 10 minutes to maintain a strong fluorescence signal inside the vessels, without leakage^51^. During the circulation period, the mice were allowed to wake up without anesthesia for better circulation. The control group did not receive bubbles + FUS treatment but was injected with the two dyes as described. Ten minutes after FITC injection, the mice were euthanized. The brains were rapidly harvested, covered in tissue freezing medium (Leica OCT cryocompound, Leica Biosystems, Nussloch GmbH, Heidelberger, Germany), and flash-frozen using liquid nitrogen. The frozen brains were then transferred to a -80ºC refrigerator for further analysis.

### Microscopy imaging and image processing

Microscopy imaging and image processing procedures were reported previously^8^. Brain sections, each with a thickness of 20 µm, were coronally sliced utilizing a cryostat microtome (CM1950, Leica Biosystems) operating at -20°C. To prevent additional extravasation of Evans Blue (EB), imaging of the brain sections occurred within 1-hour post-slicing^51^. Overview images (Fig. 2) were acquired using a motorized microscopy (Revolution, Echo, San Diego, USA). Frame images (Fig. 3 A-C) were captured using a motorized upright fluorescence microscope (BX63, Olympus, Tokyo, Japan) equipped with a x20 objective lens (UPLFLN 20X, Olympus) featuring a focal depth of 1.1 µm. Excitation wavelengths of 508 nm with an exposure time of 1150 msec were employed for the green FITC channel, while red EB channel was excited at 615 nm with an exposure time of 348 msec. Uniform imaging parameters were consistently maintained across all sections. The captured images were in dimensions of 1920 x 1200 pixels2, with each pixel corresponding to a size of 0.29 µm within the imaging plane. After image acquisition, microscopy images were processed using MATLAB (Fig. 3D). Images were split into green and red channels which marked blood vessels and EB extravasation, correspondingly. The green channel used for blood vessel segmentation into individual vessels through a modified version of the Rapid Editable Analysis of Vessel Elements Routine (REAVER) tool^52^, as discussed in detail in the automatic blood vessel segmentation section. The median radius of each segmented blood vessel was calculated according to the approach outlined by Corliss et al^52^. A binary mask based on the segmentation was applied to the red channel, effectively eliminating the interior of each blood vessel from the frame. The region of interest was determined to be 2.93 μm (equivalent to 10 pixels) and commencing at 0.59 μm (equivalent to 2 pixels) from the vessel wall around each blood vessel. The analysis of extravasation around blood vessels was performed as a function of varying distances from the vessel wall (0.3-0.9 μm) and differing extents of the perivascular area (1.5-3 μm). The intensity values within the EB channel were linearly rescaled to a [0,1] range, and the median pixel intensity within the region of interest was calculated.

### Automatic blood vessel segmentation

Automated blood vessel segmentation was performed at a single vessel resolution using a U-NET-based classifier^42^. The dataset comprised 368 green channel microscopy images of dimensions (1200x1900 pixels2). Manually labeled frames were considered as ground truth. 85% of the dataset was allocated for the training process. To enhance the quality of the training set, pixel values were initially transformed to the range [0,1], followed by linear stretching to improve contrast. Random augmentations, including variations in brightness, cropping, scaling (with a factor ranging from 0.5 to 1), rotation (up to 50°), and flipping, were introduced to mitigate overfitting. The model was trained with 100 epochs and a batch size of 8. The loss function that was used during the training was binary cross entropy combined with sigmoid:

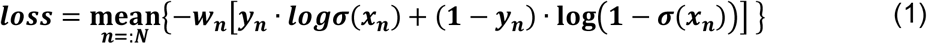

where *N* is the batch size, *y*_*n*_ is the ground truth, *x*_*n*_ is the predicted value, *w*_*n*_ is the weight and *σ* is the sigmoid function:

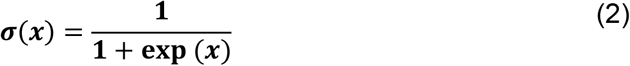

The model output was a heatmap of the same size as the input image. The output transformed the sigmoid function to yield a probability map with values in a range of [0,1]. A binary threshold of 0.3 was then applied to the results. Following the training phase, the performance of the model was evaluated by comparing its predictions on the remaining 15% of the dataset. Evaluation of the model was conducted using key metrics. First, confusion matrix was calculated^53^. Subsequently, F1 score and recall were calculated^54^:

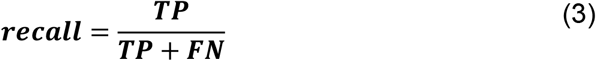

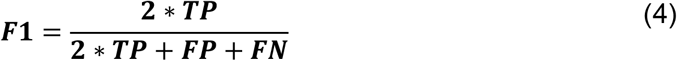

where *TP* is true positive, *FN* is false negative and *FP* is false positive. In addition, the area under the curve (AUC) of receiver operating characteristic (ROC) was calculated using scikit-learn library in Python^55^.

### Quantitative analysis

A diameter histogram was computed for each treatment group, including the control group, MB, and NB. Subsequently, these groups were subdivided by diameter in intervals of 1 µm. Since most of the blood vessels diameters were in the range of 3-4 µm, to ensure adequate sample size, blood vessels up to 8 µm in diameter were considered, ensuring a minimum of 30 blood vessels in each subgroup (Fig. 5A). Individual blood vessels underwent EB quantification (as elaborated in the image processing section), with bifurcating vessels treated as two distinct segments, one before and one after the bifurcation point. ANOVA tests were employed to compare EB intensity in the region of interest across treatment and control groups, as well as between MB and NB for each size subgroup (Fig. 5B). To assess the relative improvement of NB over MB (Fig. 5E), the ratio for each size subgroup was calculated using the following equation:

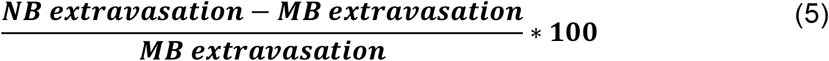

For frame analysis (Fig. 5F), for every vessel size category, the threshold for BBBD was set as a value two standard deviations greater than the average of the control group. The filtration ratio represents the proportion of open blood vessels to the total number in a given frame.

### Statistical Analysis

Statistical analysis was performed using MATLAB and GraphPad Prism. The results are presented as the mean ± 95% CI. Statistical tests are reported in the captions. P values of less than 0.05 were considered significant and were adjusted for multiple comparisons as indicated in the captions.

## Author Contributions

Conceptualization: RG, TI; Methodology: RG, LA, NR, SK; Investigation: RG, LA, NR, SK; Visualization: RG; Supervision: TI; Writing—original draft: RG, TI; Writing—review & editing: RG, TI

## Funding Sources

This work was supported by funding from Insightec Ltd., an ERC StG grant no. 101041118 (NanoBubbleBrain), the Israel Science Foundation (grant numbers 192/22 and 3450/20), the Israel Ministry of Science & Technology (grant number 101716), and the Ministry of Innovation, Science and Technology (grant number 74722).

## Data availability

Source code for the image processing algorithm will be made available upon acceptance. The datasets generated during and/or analyzed during the current study will be made available upon request.

## Declaration of competing interests

The authors have declared that no competing interest exists.

## Notes

### Competing Interest Statement

The authors have declared no competing interest.

