## Supplemental File for "Enhanced capillary delivery with nanobubble-mediated blood-brain barrier opening and advanced high resolution vascular segmentation"

**This PDF file includes:**

Supplementary Text

Fig. S1

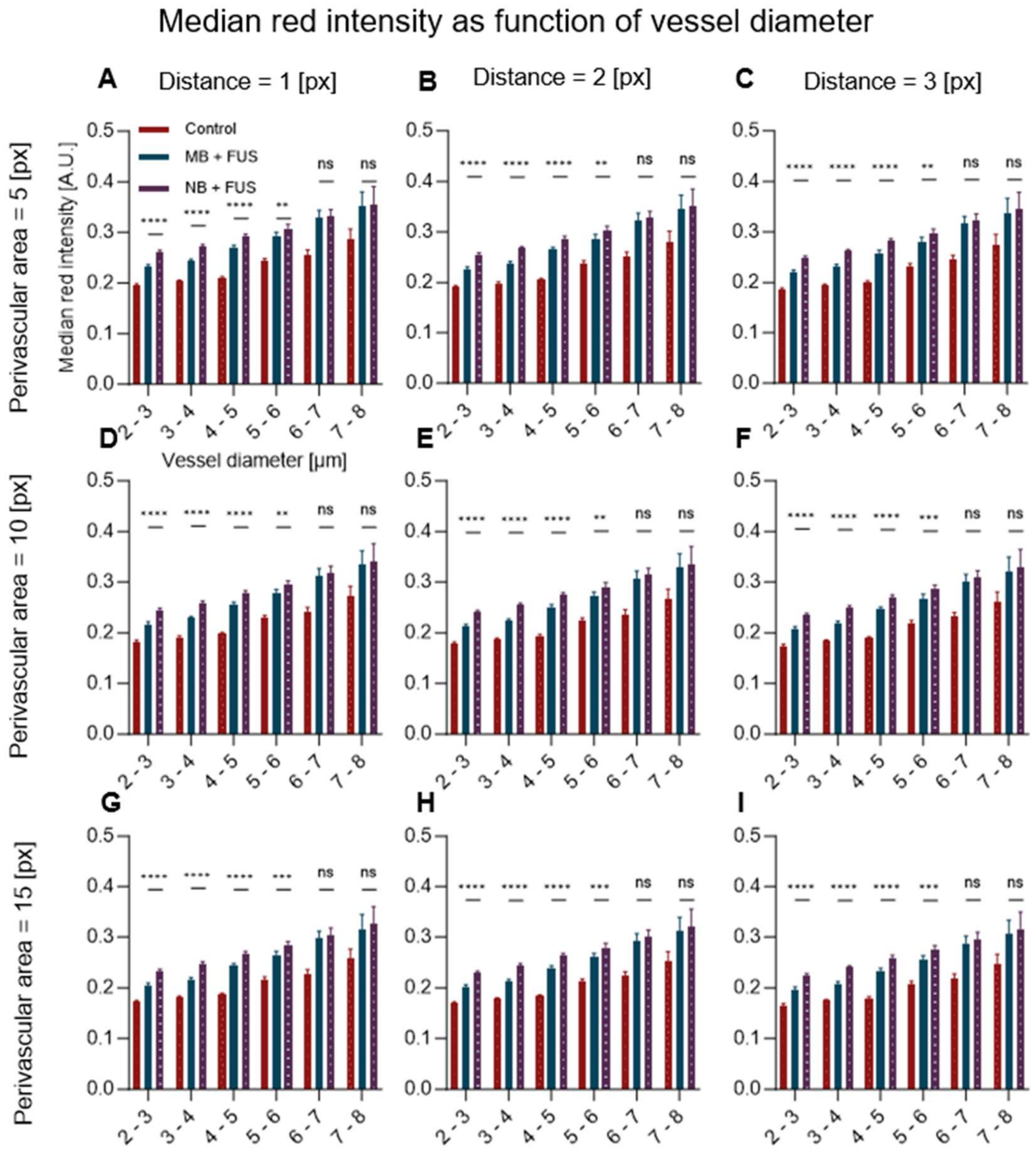


**Fig. S1. Median red intensity as function of vessel diameter and different post process parameters**. The analyzed perivascular area, denoting the length over which red extravasation is quantified, spans (A), (B), (C) 5 pixels, (D), (E), (F) 10 pixels, and (G), (H), (I) 15 pixels. The starting distance from the vessel wall, indicating where the perivascular area begins, is (A), (D), (G) 1 pixel, (B), (E), (H) 2 pixels, and (C), (F), (I) 3 pixels. 1 pixel is equivalent to 0.2929 μm. The graph is plotted as mean + 95% CI. Two-way ANOVA with Tukey correction. *$p < 0.05$, ** $p < 0.01$, ***$p < 0.001$, ****$p < 0.0001$.
